# Drugging the Medullary Thyroid Cancer Surfaceome with T Cell Engagers Targeting CEA, GFRA4 and DLL3 as Monotherapy and In Combination with Tyrosine Kinase Inhibitors

**DOI:** 10.1101/2025.08.05.668474

**Authors:** Tim Andrew Erickson

## Abstract

Medullary thyroid cancer (MTC) is a rare form of thyroid cancer, and the definitive treatment is surgical resection. For patients who are not cured by surgery or for patients who present with distant metastatic disease, no curative therapy exists. T cell engagers (TCEs) are an emerging class of biologics that simultaneously bind to tumor cell surface antigens and to the CD3 domain of T cells, thereby redirecting endogenous T cells to lyse tumor cells. TCEs can mediate durable tumor regression, including complete responses, and are now approved to treat multiple cancer types outside of MTC. RNASeq analysis of 30 MTC tumors, cross-referencing with public databases and an extensive literature review were used to identify CEA, DLL3 and GFRA4 as promising tumor antigens to target with TCEs. Expression of these antigens was then validated on MTC cell lines. Herein, I describe the discovery, application and development of MTC-targeted TCEs (MTCEs) recognizing the MTC tumor antigens CEA, DLL3 and GFRA4. *In vitro*, MTCEs mediate potent target-dependent and T cell-dependent cytotoxicity against MTC cell lines at concentrations as low as 10 ng/mL. MTCEs also induce target-dependent IFNγ secretion from T cells, with no measurable IFNγ secretion observed in the presence of antigen-negative cell lines. Further, MTCEs are functionally compatible with the FDA-approved tyrosine kinase inhibitors selpercatinib and cabozantinib, and combination therapy numerically enhances cytotoxicity. These preclinical data provide strong rationale for continued development of MTCEs, which may one day revolutionize the treatment of metastatic MTC.

## Introduction

Medullary thyroid cancer (MTC) is a relatively rare thyroid cancer that originates from the neuroendocrine parafollicular C-cells of the thyroid [1]. While localized MTC can sometimes be cured surgically, there is presently no cure for metastatic MTC [2]. While high-dose stereotactic body radiotherapy (SBRT) can provide local control of individual tumors [3] and Y-90 radioembolization can mediate durable control of liver metastases [4], there is no highly effective adjuvant therapy for MTC capable of eliminating all sites of metastatic disease. Unlike differentiated thyroid cancers, MTC does not concentrate radioactive iodine [5] and MTC is generally resistant to cytotoxic chemotherapy, as evidenced by low response rates and the transient nature of responses [6].

In the setting of progressive metastatic MTC, the daily oral tyrosine kinase inhibitors (selpercatinib, cabozantinib and vandetanib) can prolong survival for MTC patients harboring RET mutations, at the expense of side effects and cumulative toxicities [7–9]. In particular, selpercatinib treatment resulted in high response rates and a robust multi-year overall survival (OS) benefit in RET-mutant MTC [9]. However, no randomized trial to date has demonstrated any OS benefit in RET wild-type patients [7]. Thus, there is a medical need to move the field forward and explore additional therapeutic strategies with the moonshot aim of curing metastatic MTC.

T cell engagers (TCEs) are an emerging class of biologics that simultaneously bind to tumor-associated surface antigens and to CD3 expressed on T cells, forming an immunological synapse, resulting in T cell activation and tumor cell lysis [10]. During the last decade, multiple TCEs targeting CD19, CD20 and BCMA have been approved to treat hematological malignancies [11]. The 2024 FDA approval of the anti-DLL3 TCE tarlatamab in small cell lung cancer (SCLC) marked the first approval of a TCE in any solid tumor type [12]. In recent clinical trials, TCEs targeting STEAP1 and PSMA have shown robust antitumor activity in treatment-refractory prostate cancer, including durable responses and 99% PSA declines [13,14]. Further, TCEs targeting CEACAM5 (CEA) [15] and EGFR [16] have mediated objective responses in clinical trials in GI malignancies.

The successful application of TCEs in MTC would represent a new treatment paradigm, enabling the direct eradication of MTC tumor cells by endogenous T cells, in a tumor type which has historically proven resistant to immunotherapies, including immune checkpoint inhibitors and tumor-antigen vaccines [17,18]. While TCEs have been evaluated in multiple tumor types, PubMed and Google Scholar searches did not yield any publications evaluating TCEs in MTC, to date. Herein, the discovery, development and *in vitro* preclinical activity of TCEs targeting the MTC tumor antigens CEA, DLL3 and GFRA4 is described. These MTC-targeting TCEs are termed MTCEs. The identification of CEA, DLL3 and GFRA4 as addressable target antigens based on patient data is first discussed followed by target validation on MTC tumor cells lines using flow cytometry. Then, the development of MTCEs is described, and their independent binding to both T cells and target antigens is quantified using flow cytometry. Lastly, the *in vitro* activity of MTCEs alone and in combination with the FDA-approved kinase inhibitors cabozantinib and selpercatinib is characterized through live-cell imaging, cytotoxicity assays and measurement of MTCE-mediated IFNγ secretion in tumor cell: T cell co-cultures.

## RESULTS

### Identification of Target Antigens for MTCEs

30 MTC patient tumors (26 primary; 4 tumor infiltrated lymph nodes (TILN)) were analyzed by bulk RNA sequencing, as previously described [19]. An analysis of the RNASeq dataset in conjunction with Human Protein Atlas curation and a literature review was used to identify target antigens for MTCEs. To conduct the analysis, the MTC patient dataset was truncated to exclude genes coding for intracellular proteins, leaving a set of 3,412 genes which code for the extracellular protein surfaceome [20]. MTC tumor surfaceome genes were then ranked by median expression level. Next, to identify genes specifically upregulated in MTC tumors, and to exclude genes with global elevated gene expression, the surfaceome of the TRON cell line database [21] was analyzed by dividing the expression value of each gene in the MTC TT cell line by the corresponding median expression value of all other 1,081 non-MTC cell lines in the TRON database (TT cell line/TRON cell line median). Based on the differential gene expression analysis, the top 40 relatively overexpressed TT surfaceome genes were selected for further analysis. Then, analysis was undertaken to rank genes based on both high differential expression and high expression in the MTC patient dataset, noting that this approach filtered out non-MTC genes present in the MTC dataset, such as TPO and TSHR due to the presence of residual thyroid tissue.

The product of differential gene expression and median MTC patient expression was computed and candidate genes were ranked accordingly (Table 1). Lastly, each gene was annotated according to normal tissue expression in the Human Protein Atlas [22], in order to exclude genes with broad expression in normal tissues and simultaneous high expression in MTC, such as RET. An exception was made for CEACAM5 (CEA) because of the reported aberrant polarization (basal membrane expression) in tumor cells, which should theoretically limit on-target, off-tumor toxicity as T cells can not readily access the lumen-facing apical membrane, where CEA exhibits restricted expression in normal epithelial tissues [23]. Per the aforementioned analysis, CEA, GFRA4, DLL3, GHSR, APLP1, PCDHA5 and PCDHA8 were identified as candidate target genes (Figure 1), as they showed high relative expression and differential expression in MTC and restricted normal tissue expression.

**Figure 1:**
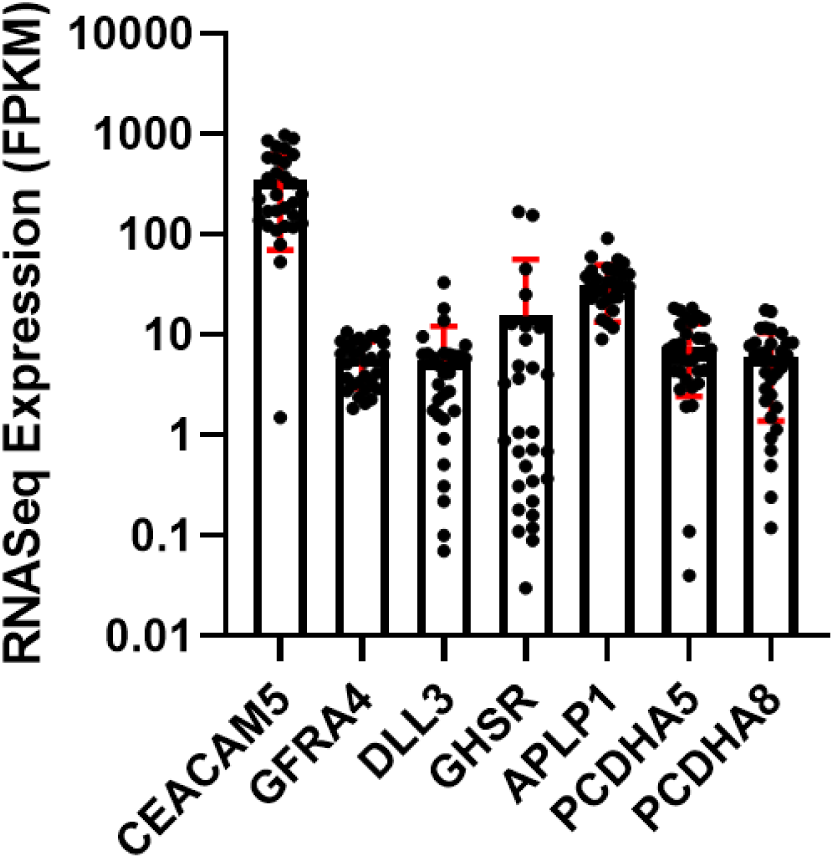
RNA expression in FPKM of candidate target antigens from a dataset comprised of 30 MTC patient tumors (26 primary tumor and 4 lymph-node infiltrated tumors). Paraffin-embedded tumor tissue blocks were used as input material.

**Table 1:**
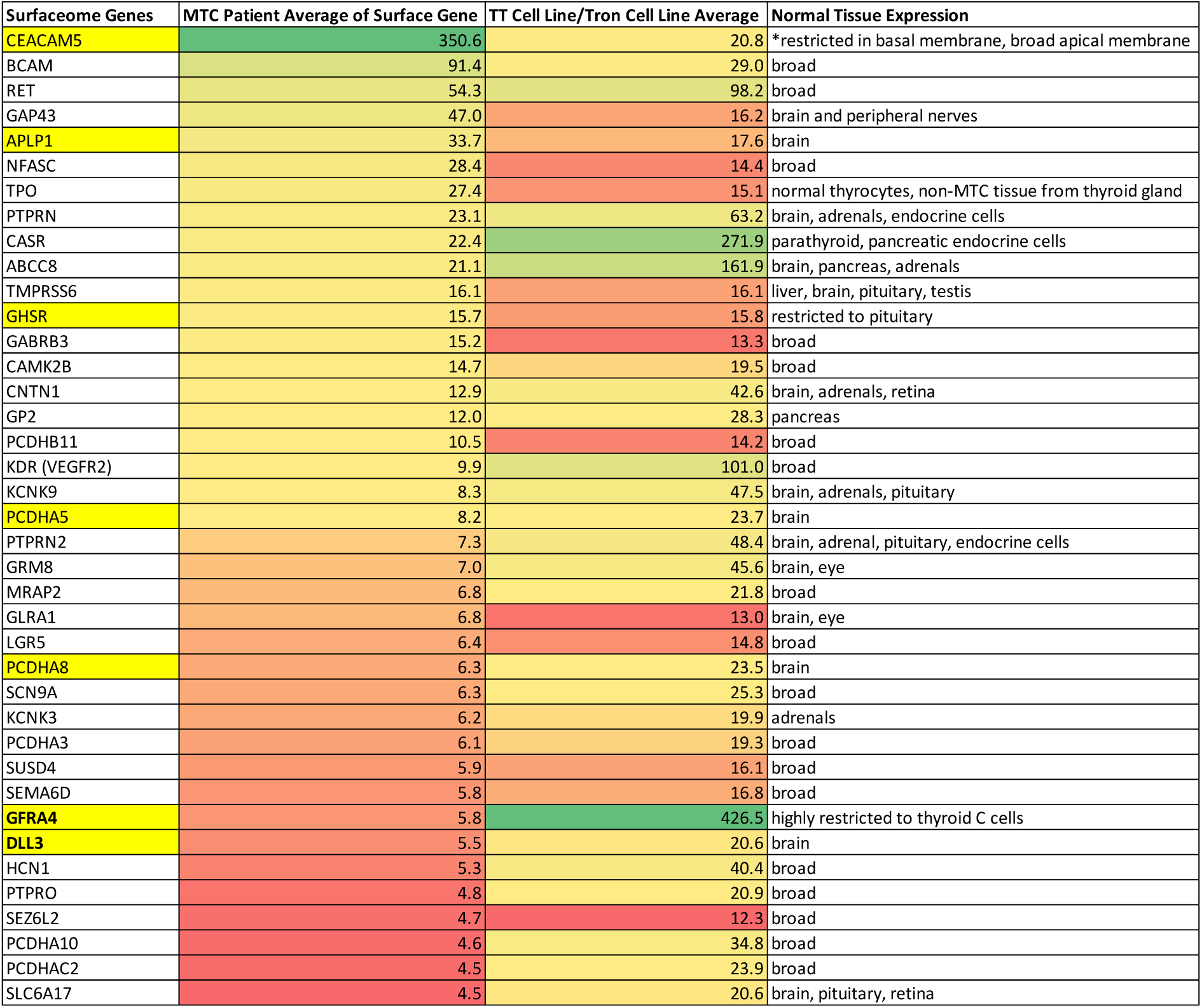
MTC tumor RNASeq analysis, differential gene expression analysis and normal tissue expression analysis used to identify MTCE target antigens.

As indicated in Figure 1, the well-established MTC tumor marker CEA [24] showed the highest expression, while the RET co-receptor GFRA4 (GDNF family receptor alpha-4) showed the highest differential gene expression in the TRON cell line database, in agreement with the previous reports indicating that GFRA4 expression is reported to be highly restricted to thyroid C cells and MTC [25]. Notably, both anti-CEA and anti-GFRA4 chimeric antigen receptor (CAR) T cells have demonstrated robust preclinical activity in MTC [26,27], and a clinical trial is currently evaluating anti-GFRA4 CAR-T cells in MTC patients [28]. Further, a clinical trial of the anti-CEA TCE cibisatamab in colon cancer mediated objective responses without dose limiting toxicities [15].

DLL3, GHSR, APLP1, PCDHA5 and PCDHA8, all show normal tissue expression restricted to the regions of the brain, which may preclude these antigens from being targeted due to the potential for neural toxicity. Surprisingly, DLL3 has proven to be a safe target [29], potentially due to aberrant extracellular trafficking in tumor cells [30]. Recent reports have indicated that DLL3 is abundantly expressed at both RNA and protein levels in aggressive metastatic MTC, which often harbors a transcriptomic signature (DLL3, SOX2, ASCL1) akin to SCLC [31]. Further, DLL3 has been detected in up to 100% of tumor cells in lymph node metastases by IHC [32]. Based on the totality of data, including high expression, relative overexpression, validated expression and indications of safety, CEA, DLL3 and GFRA4 were selected as target antigens for MTCEs. Of the remaining 4 candidate genes in the curated list, APLP1 shows the highest and most uniform expression and may be a suitable target although it was not selected for further development due to a lack of protein expression data in the literature and documented RNA expression in the human brain.

### Expression of CEA, DLL3 and GFRA4 on MTC Cell Lines and the SCLC Cell Line SHP-77

For proof-of-concept studies, flow cytometry was used to measure the expression of CEA, DLL3 and GFRA4 in the TT and MZ-CRC-1 (MZ) MTC cell lines and non-MTC cell lines (SHP-77, LNCaP, Raji, SKOV3, CHO-wt). The SCLC cell line SHP-77 shares a neuroendocrine origin with MTC and is known to express DLL3 and CEA [21], whereas the other cell lines are negative for all three targets. As shown in Figure 2, TT cells highly and broadly express CEA, DLL3 and GFRA4 with nearly 100% of cells expressing all three targets. In contrast, MZ cells show almost no detectable expression of GFRA4, moderate CEA (30%) expression and broad DLL3 expression. SHP-77 cells were found to broadly express DLL3 [33], and also show moderate CEA expression. SHP-77 cells are not reported to express GFRA4 at the RNA level [21], whereas low-level RNA expression (100-fold less than TT cells) has been reported for MZ cells [27]. The detection of very dim populations can be challenging using multiparameter flow cytometry, due to cytometer drift and slight non-linearities in photodetector response, which limit the absolute accuracy of linear compensation matrices. As such, single antibody staining is more robust for the detection of dim protein populations. A subsequent single staining experiment was conducted to analyze GFRA4 expression on the SHP-77 and MZ cells using PE-labeled anti-GFRA4. The results (Figure S1) demonstrate no detectable expression of GFRA4 on SHP-77 cells relative to isotype control. Conversely, very dim expression of GFRA4 is detectable on MZ cells. CEA, DLL3 and GFRA4 were not detectable by flow on any of the other cell lines (data not shown).

**Figure 2:**
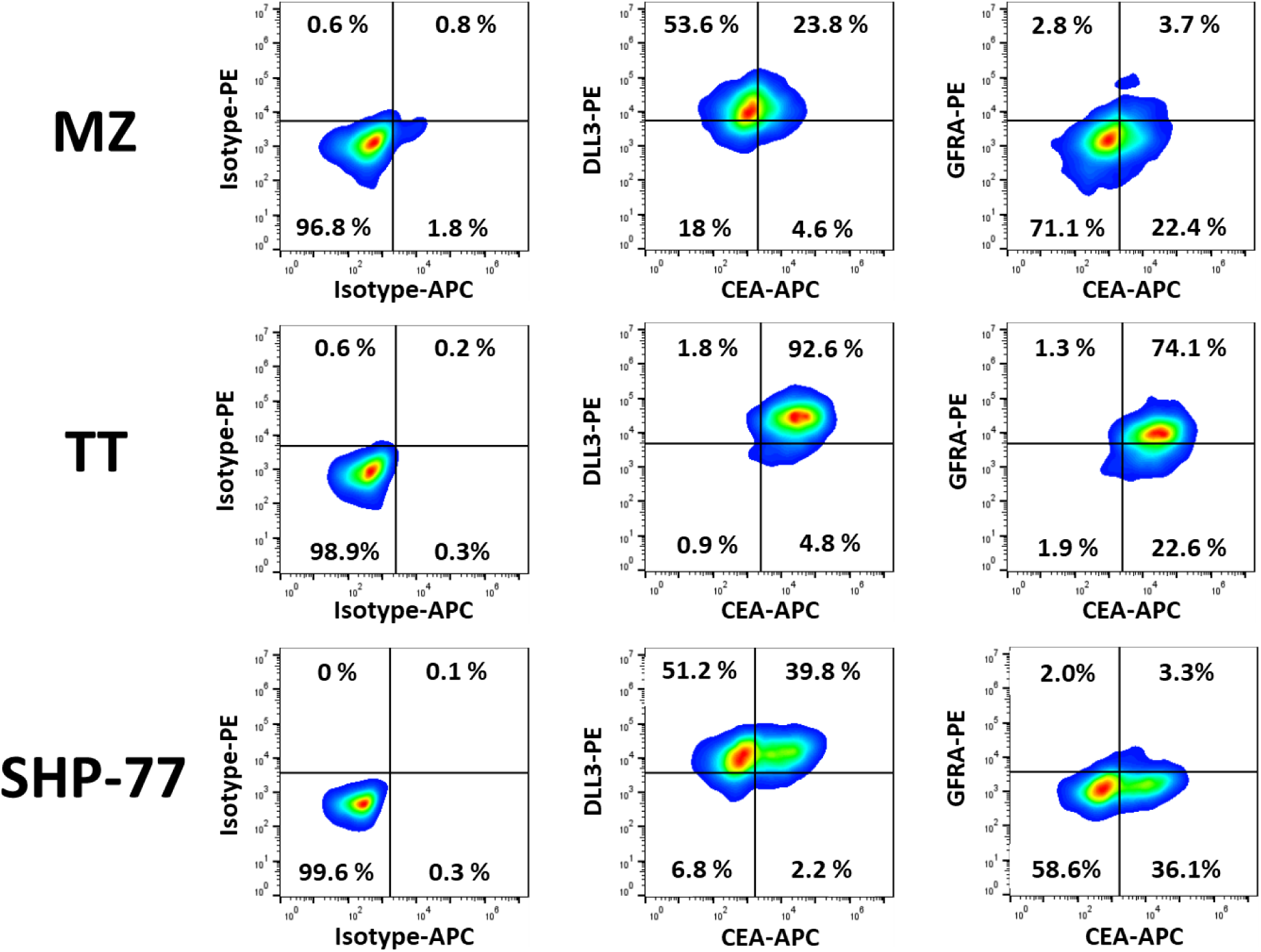
Expression of CEA, DLL3 and GFRA4 on MZ (MTC), TT (MTC) and SHP-77 (SCLC) cell lines as measured by flow cytometry. TT cells show high expression of all three antigens, whereas MZ and SHP-77 cells do not express significant amounts of GFRA4.

### Engineering and Synthesis of MTCEs

Humanized monoclonal antibodies recognizing CEA and GFRA4 were generated through murine immunization, followed by AI-based affinity maturation and humanization by grafting the light chain and heavy chain CDRs onto homologous human antibody frameworks. Single chain variable fragments (scFv) targeting CEA and GFRA4 were then generated by linking the heavy and light chain variable regions of each monoclonal antibody using a (GGGGS)_3_ linker. In order to generate MTCEs targeting CEA (anti-CEA) and GFRA4 (anti-GFRA4), each respective scFv was in turn linked to a proprietary anti-CD3 scFv/half-life extension domain. Specifically, the anti-CD3 domain was derived from a humanized scFv of the SP34 anti-CD3 antibody [34], and the half-life extension domain was derived from an Fc-silenced constant region of human IgG4. For proof-of-concept studies, tarlatamab was selected as the candidate anti-DLL3 MTCE because it is already FDA approved in SCLC and could be readily used to treat MTC patients off-label.

Anti-CEA and anti-GFRA4 MTCEs were expressed via transient transfection of CHO cells with plasmids encoding each MTCE followed by Protein A and size exclusion chromatography purification. Purity exceeded 93% as measured by SDS-PAGE and SEC-HPLC with yields of approximately 300 mg/L. The anti-DLL3 MTCE tarlatamab, cibisatamab, and anti-CLDN18.2 (gresonitamab) were all purchased directly from a commercial vendor (MedChemExpress). All MTCEs were tested for endotoxin and confirmed to have levels of <0.1 EU/mg. The general structure of each MTCE, expressed as a continuous single-chain protein, is depicted in Figure 3.

**Figure 3:**
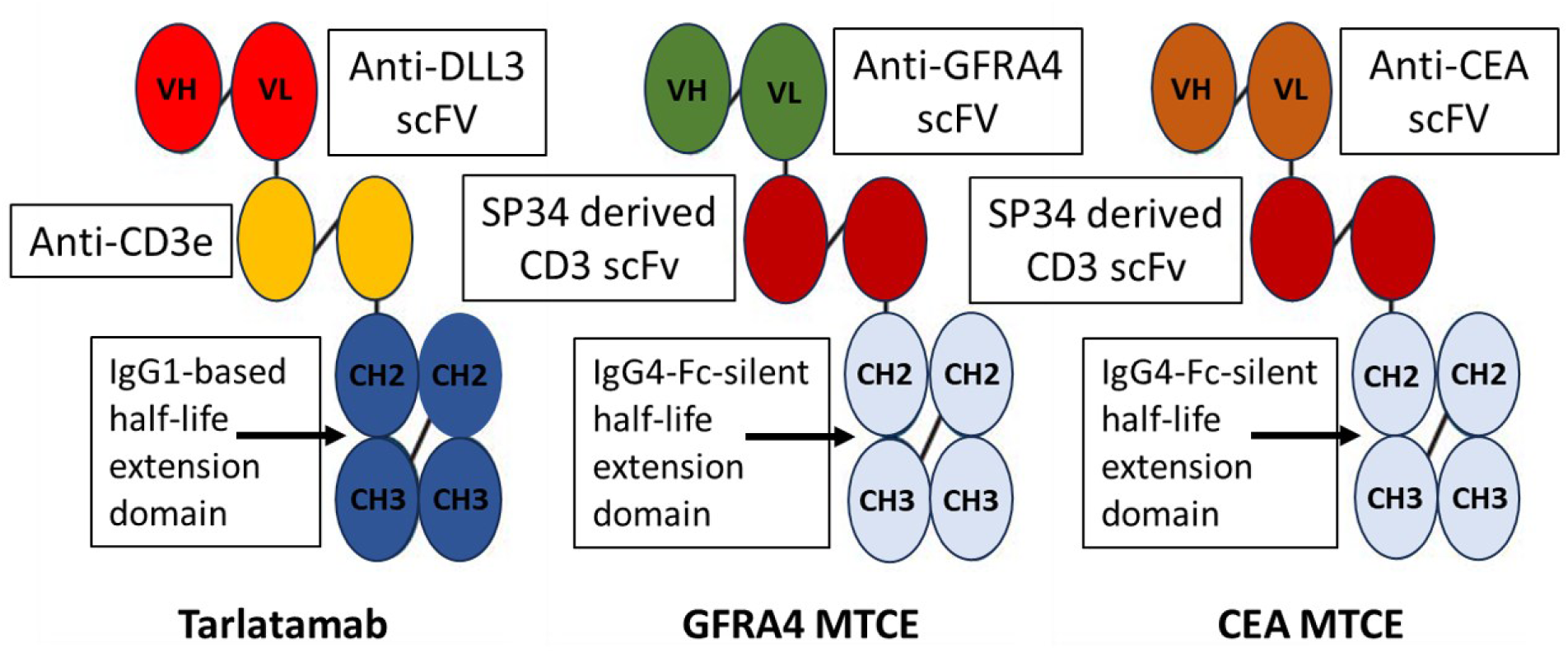
Generalized structure of tarlatamab (anti-DLL3) and anti-CEA and anti-GFRA4 MTCEs. Each TCE is comprised of an antigen binding domain, a CD3 binding domain and a Fc-based half-life extension domain.

### Functional Binding of MTCEs to T Cells and Target Antigens

Flow cytometry-based assays were used to characterize the functional binding of MTCEs to T cells and tumor antigens. For T cell binding, T cells were first isolated from peripheral blood PBMCs using negative selection. Purity was checked by flow and found to exceed 97%. Purified T cells were then incubated with varying concentrations (100 pg/mL to 10 µg/mL) of MTCEs for 20 minutes at room temperature, then washed twice and counterstained with FITC-conjugated CD8 mAb, PerCp-conjugated CD4 mAb and 1 µg/mL of PE-conjugated goat anti-human IgG antibody fragment, which binds to the Fc-domain of the MTCEs. Cells were washed to remove unbound antibodies, and flow data were acquired on a BD Accuri cytometer. Background subtraction at each MTCE concentration was used in the final computation by subtracting the Median Fluorescence Intensity of the isotype control antibody from the measured MTCE Median Fluorescence Intensity at the same concentration (x) such that MFI_MTCE,x_ = (MFI_measured MTCE,x_-MFI_isotype,x_). As demonstrated in Figure 4, all MTCEs bind to CD8 T cells with similar affinity, and binding increases monotonically with MTCE concentration with detectable binding observed at concentrations as low as 10 ng/mL. Similar binding for each MTCE was observed for CD4 T cells, whereas no binding was observed for CD20+ B cells, CD14+ monocytes, or CD16+CD56+ NK cells (data not shown).

**Figure 4:**
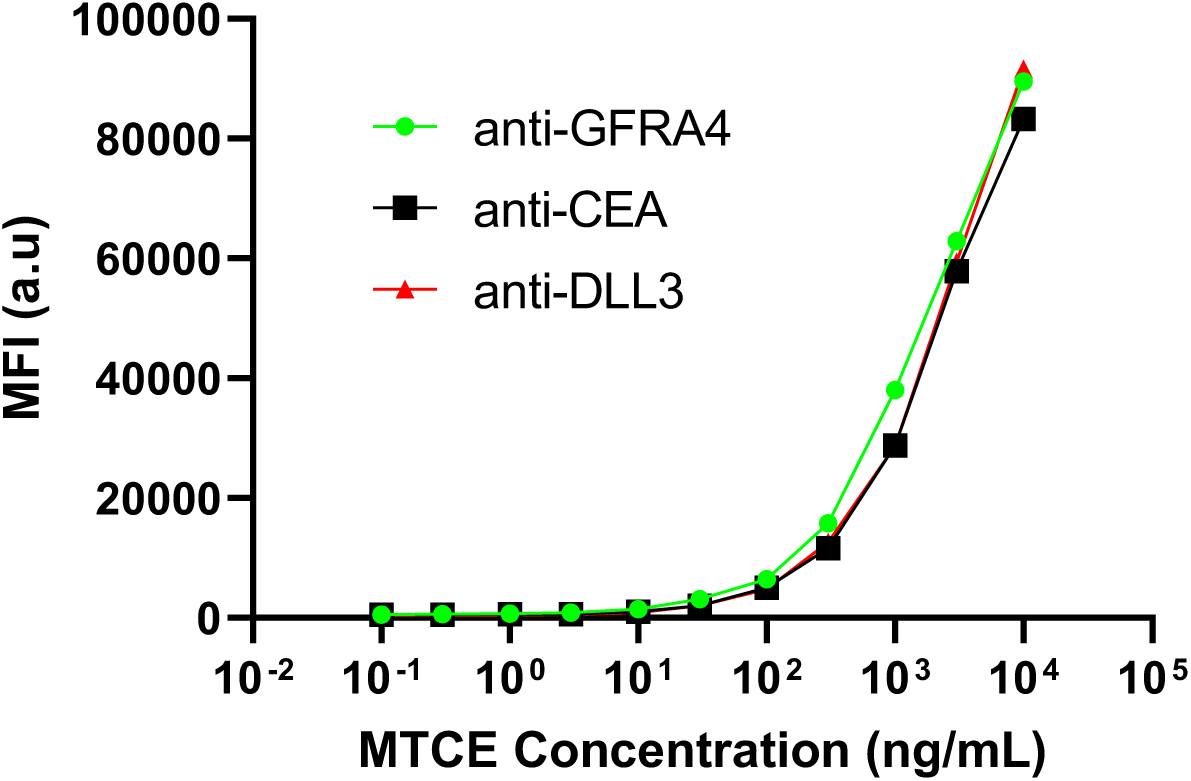
Binding of MTCEs to CD8 T Cells. Purified CD8 T cells were incubated with varying concentrations of MTCEs for 20 minutes, then washed twice in ice-cold FACS buffer and counterstained using anti-human Fc PE antibody, then washed twice again with ice-cold FACS buffer to remove unbound anti-human Fc antibody prior to flow cytometry analysis. Binding was quantified as Median Fluorescence Intensity (MFI) on the PE-channel, which assuming photodetector linearity, is proportional to the number of bound MTCE molecules. Measurable binding was observed at just 10 ng/mL and binding increases monotonically up to 10 µg/mL.

For tumor antigen binding analysis, CHO cells were transduced with lentiviral vectors encoding human CEA (NP_004354.3), DLL3 (NP_058637.1) or GFRA4 (NP_665705.1). Limiting dilution was used to select clonal populations with high and homogenous expression for each antigen. Target antigen surface expression was confirmed on CHO cells using flow cytometry (Figure S2). CHO-wt, CHO-CEA, CHO-DLL3 and CHO-GFRA4 cells were then incubated with varying concentrations (100 pg/mL to 10 µg/mL) of MTCEs for 20 minutes at room temperature, washed twice with ice-cold FACS buffer and then counterstained with PE-conjugated goat anti-human IgG, washed twice in ice-cold FACS buffer and then analyzed on a BD Accuri flow cytometer. Background correction was employed, as previously described. No detectable binding was observed for any MTCEs to CHO-wt cells and no cross-reactive binding was observed for MTCEs against non-target transduced CHO cells, i.e. binding of anti-CEA MTCE to GFRA4-transduced CHO cells. Figure 5 shows the on-target binding data for each MTCE, which was normalized to account for differences in target-antigen expression levels on CHO cells. Whereas anti-DLL3 and anti-CEA MTCEs showed similar binding affinity, with binding clearly detectable at concentrations as low as 3 ng/mL, there was an approximate log-shift in binding affinity (EC_50_) for anti-GFRA4 MTCE where minimal binding was observed at a concentration of 30 ng/mL.

**Figure 5:**
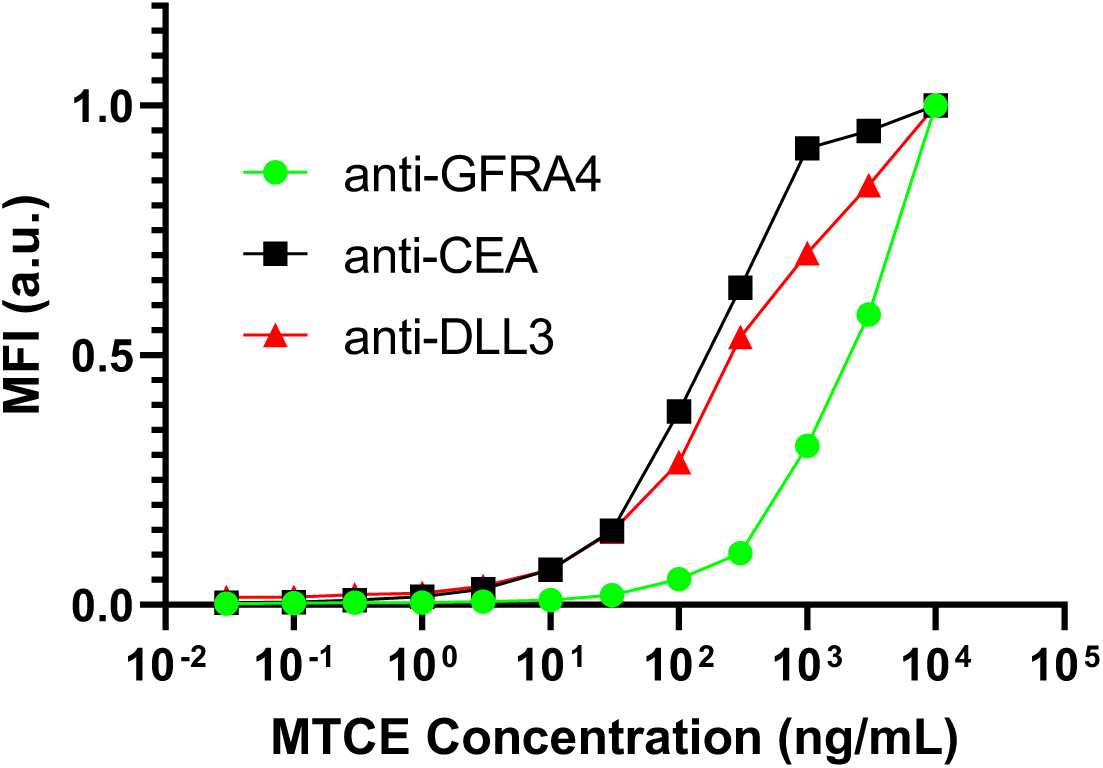
Normalized binding of MTCEs to target-transduced CHO cells. The data shows binding of anti-GFRA4 MTCE to CHO-GFRA4 (green circles), binding of anti-CEA MTCE to CHO-CEA (black squares) and binding of anti-DLL3 MTCE to CHO-DLL3 (red triangles). The data is reported as normalized Median Fluorescence Intensity on the PE-channel, which assuming photodetector linearity, is proportional to the number of bound MTCE molecules.

### MTCEs Mediate Cytotoxicity Against MTC Cell Lines

MTCEs were capable of mediating potent cytotoxicity against MTC cells in the presence of T cells. The XTT cytotoxicity assay and endpoint live-cell imaging were used to assess MTCE-dependent and T-cell dependent cytotoxicity against MTC cell lines *in vitro.* Briefly, the XTT assay is based on the conversion of the tetrazolium salt XTT into an orange formazan product by metabolically active adherent tumor cells, and the colorimetric shift is quantified by absorption spectroscopy. Separate experiments were conducted to characterize the influence of both MTCE concentration and E:T ratio on cytotoxicity. Each experimental condition was run in triplicate using T cells from the author.

In the MTCE titration experiment, the E:T ratio was fixed at 1:1. Briefly, 15,000 GFP-transduced MTC cells were plated overnight in each well of a 96-well flat bottom plate, and the following day, 15,000 T cells were added to each well along with varying concentrations of each MTCE. Cells were co-cultured for 48 hours, and then endpoint live-cell imaging was conducted on a fluorescence microscope before performing the XTT assay. The results in Figure 6 demonstrate how each MTCE mediated cytotoxicity against MZ and TT cells in a concentration-dependent manner. MTCEs targeting DLL3 and CEA exhibited potent cytotoxicity against both cell lines, whereas anti-GFRA4 was only effective against the TT cell line, consistent with the lack of expression of GFRA4 on the MZ cells. In general, anti-DLL3 (tarlatamab) mediated the broadest and most potent cytotoxicity across both cell lines with an approximate EC50 of ∼2 ng/mL for TT cells and 0.2 ng/mL for MZ cells. Anti-CEA MTCE induced potent cytotoxicity in TT cells with an EC50 similar to tarlatamab, but showed reduced cytotoxicity in the MZ cell line, consistent with the reduced expression of CEA in the MZ cell line. While anti-GFRA4 induced potent cytotoxicity against TT cells at high concentrations, it had the highest EC50 value (EC50_TT_ ∼ 20 ng/mL and EC50_MZ_ ∼ 10 µg/mL, which may be a function of reduced binding affinity compared to anti-CEA and anti-DLL3 (Figure 5) or potentially lower GFRA4 protein expression relative to DLL3 and CEA.

**Figure 6:**
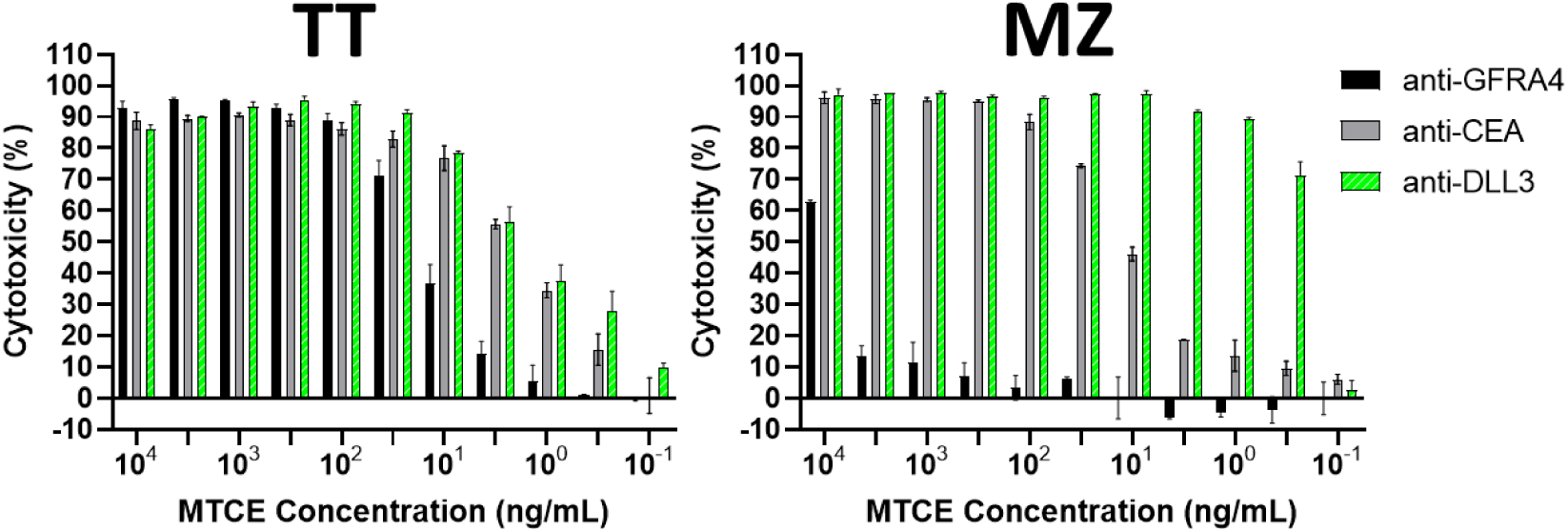
Concentration-dependent cytotoxicity of MTCEs against the MZ and TT MTC cell lines at a 1:1 E:T ratio. Tumor cells were co-cultured with T cells for 48 hours in the presence of MTCEs as indicated, and then cytotoxicity was measured using the XTT assay.

For the T cell titration experiment, 15,000 tumor cells were again plated overnight in 96-well plates, and the following day T cells were added at E:T ratios ranging from 1:1 to 1:32 in the presence of 1 µg/mL of each MTCE. Both media only and the anti-CLDN18.2 TCE gresonitamab were used as negative controls, as CLDN18.2 is not expressed on MTC cell lines. After 24 hours, live-cell imaging was performed on an inverted fluorescence microscope for all TCEs at the 1:2 E:T ratio condition, in order to visually confirm cytotoxicity. As shown in Figure 7, all three MTCEs mediated potent cytotoxicity against TT cells as evidenced by reduced GFP surface area and clumping of GFP+ tumor cells. By contrast, T cells alone or T cells in the presence of gresonitamab did not mediate appreciable cytotoxicity, as the appearance is similar to TT cells in media only. For MZ cells, only tarlatamab induced high, visually discernible cytotoxicity, whereas anti-CEA cytotoxicity was discernible, but much less pronounced, and neither anti-GFRA4, gresonitamab nor T cells alone mediated visually detectable cytotoxicity.

**Figure 7:**
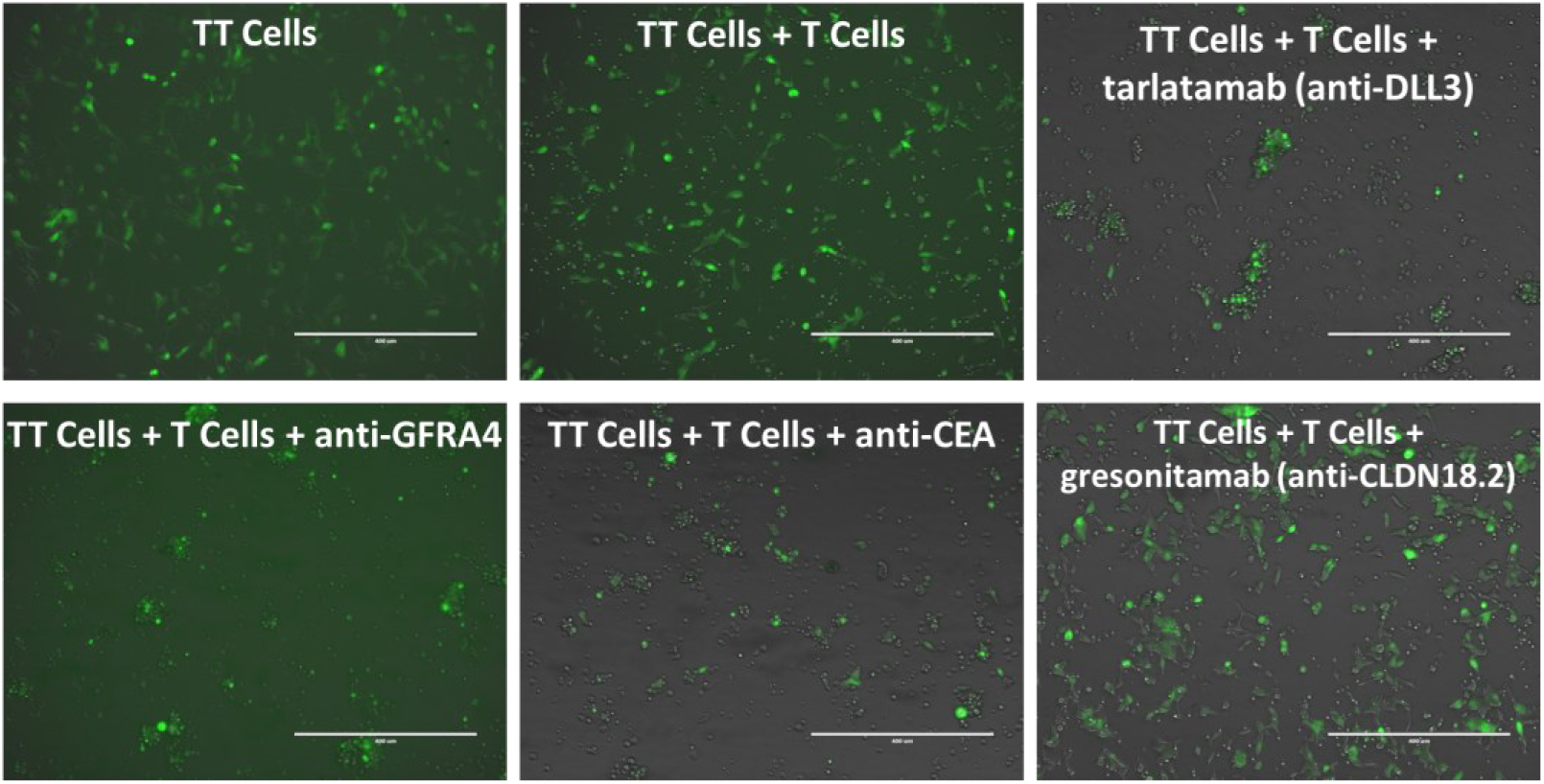
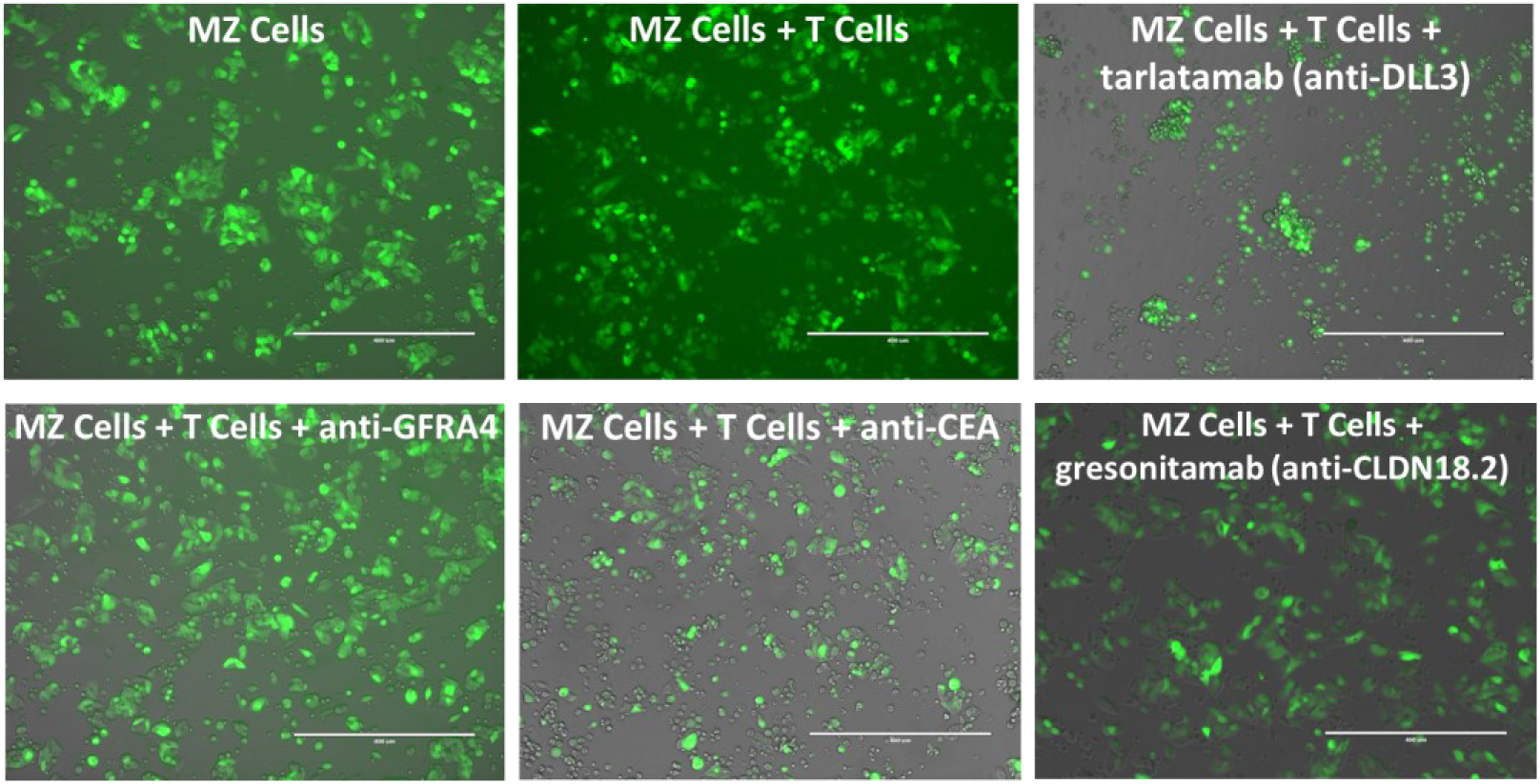
Live-cell imaging reveals lysis of MTC cells mediated by T cells in the presence of MTCEs. All three MTCEs mediated observable cytotoxicity against the TT cell lines, whereas only anti-DLL3 and anti-CEA mediated observable clearance of MZ cells. The scale bar is 400 microns.

XTT assay results (Figure 8) demonstrate that MTCEs exhibited T cell-dependent cytotoxicity, with cytotoxicity beginning to fall off at E:T ratios of 1:4 and lower. Notably, tarlatamab demonstrated the highest and most consistent cytotoxicity at the lowest E:T ratio (1:32), and no detectable cytotoxicity was observed in the absence of MTCEs, nor in the absence of T cells or with gresonitamab at any E:T ratio (data not shown).

**Figure 8:**
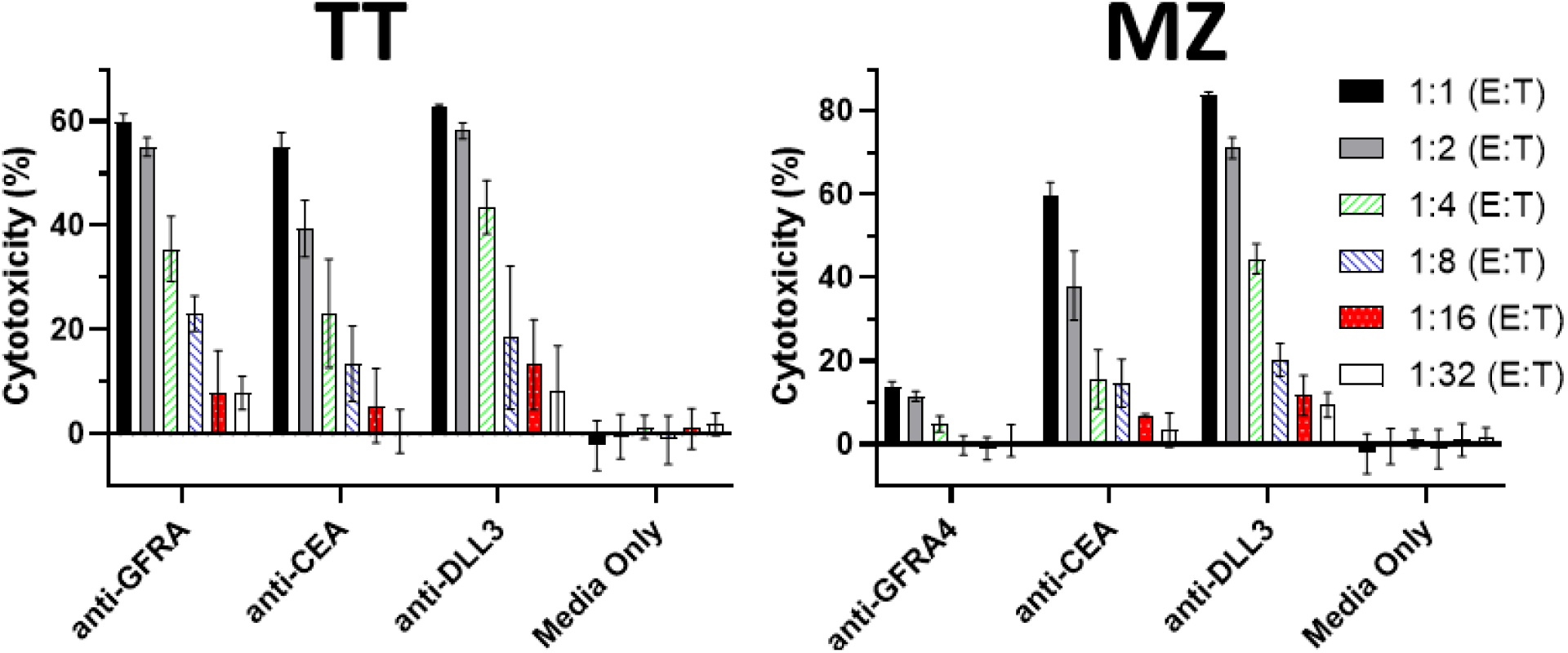
MTCE-mediated cytotoxicity requires T cells and depends on the effector to target (E:T) ratio.

### MTCEs Mediate Cytotoxicity in a Target-Dependent Manner

To determine the impact of target antigen expression on cytotoxicity, a 48-hour XTT co-culture assay was conducted with 10 µg/mL of each MTCE at 1:1 E:T ratio (15,000 T cells, 15,000 tumor cells in a 96-well plate) for 48-hours. Antigen-positive (MZ (CEA+, DLL3+), TT (triple positive), SHP-77 (CEA+, DLL3+), CHO (single positive)) and antigen-negative (LNCaP, SKOV3, Raji, CHO-wt) cell lines were used in the assay. Gresonitamab was used as a negative control, and the CEA-reactive TCE cibisatamab was used as a positive control for CEA+ cell lines and a negative control for CEA-cell lines. As shown in Figure 9, MTCEs exhibited cytotoxicity in an antigen-dependent manner. Notably, MTCEs only mediated cytotoxicity against CHO cells transduced to express their target antigen. Further, while both anti-CEA and cibisatamab mediated cytotoxicity against CEA+ cells lines, anti-CEA MTCE exhibited more potent cytotoxicity than cibisatamab in MZ cells, TT cells, SHP-77 cells and CHO-CEA cells. Critically, MTCEs did not mediate detectable cytotoxicity against antigen-negative cell lines at the highest E:T ratio and highest MTCE concentration tested, and no appreciable cytotoxicity was observed for cells exposed to gresonitamab.

**Figure 9:**
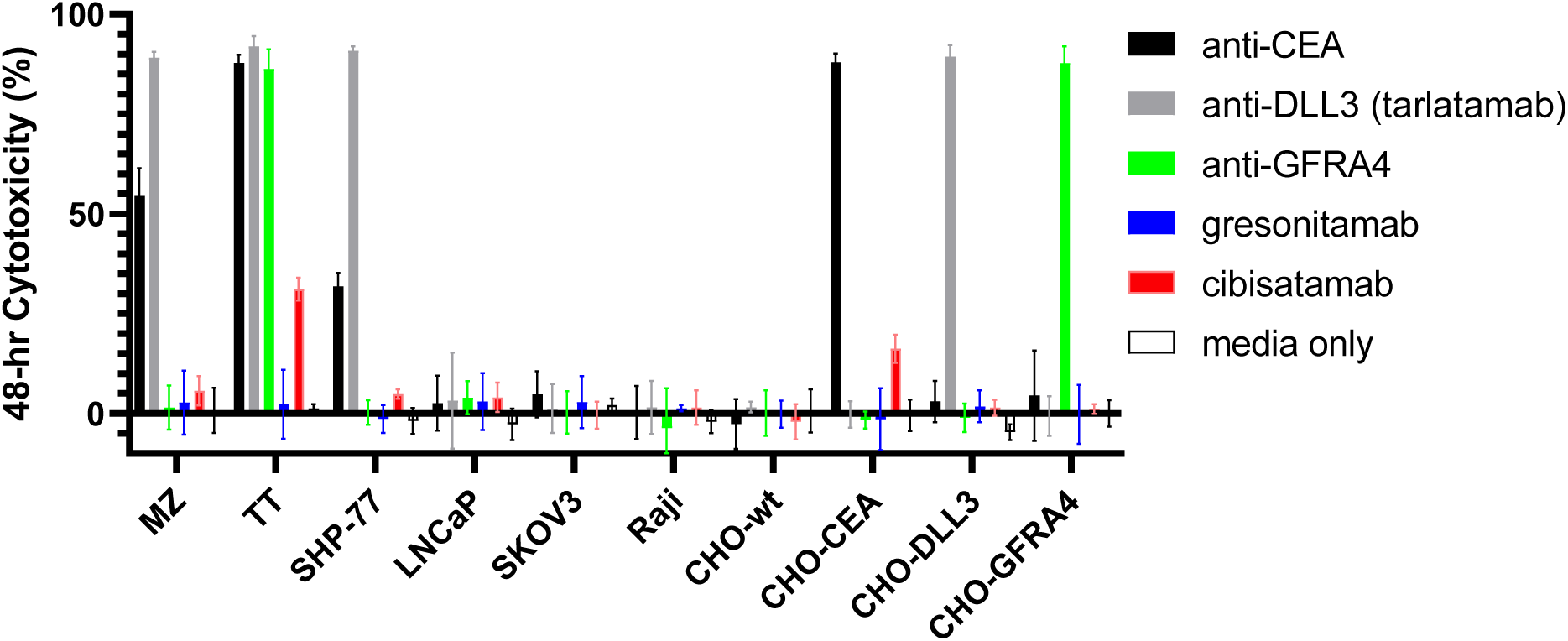
MTCEs mediate cytotoxicity in a target-dependent manner with no off-target cytotoxicity observed in target antigen-negative cell lines.

### Combining MTCEs with Tyrosine Kinase Inhibitors (TKIs) Increases Cytotoxicity

The RET-specific tyrosine kinase inhibitor selpercatinib and the multikinase inhibitor cabozantinib are both approved to treat metastatic MTC, and intriguingly both have been reported to enhance immunotherapy by reducing immunosuppression in the tumor microenvironment (TME) by upregulating MHC Class I, increasing T cell infiltration, and reducing the presence and function of immunosuppressive cellular subsets [35–37]. As TKIs may further potentiate the activity of MTCEs *in vivo* through several mechanisms, both cytotoxicity and IFNγ secretion assays were used to assess the *in vitro* activity and compatibility of MTCEs with TKIs at clinically relevant concentrations of 5 µM for selpercatinib and 3 µM for cabozantinib [38,39]. To model a patient undergoing continuous TKI treatment followed by MTCE therapy, 15,000 MZ and TT cells were plated overnight and then exposed to each TKIs (or media only) for 2 days, after which the media was refreshed with the same concentration of TKIs and then 7,500 T cells were added with the 1 µg/mL of each MTCE. The co-culture was then carried out for 48 hours, after which IFNγ secretion and cytotoxicity were measured. As shown in Figure 10, the addition of selpercatinib or cabozantinib generally decreased IFNγ secretion, but did not abrogate it entirely. This phenomenon may be a function of reduced tumor cell volume, which in turn my reduce the total surface area available for T cell binding and activation. Interestingly, the reduction in IFNγ did not lead to reduced cytotoxicity. Rather, trends toward increased cytotoxicity were observed for all MTCEs in the presence of selpercatinib and cabozantinib. These data indicate that TKIs are likely functionally compatible with MTCEs, as they do not hinder cytotoxicity, and that TKIs may reduce unwanted side effects, such as cytokine release by reducing tumor surface area. Notably, while anti-GFRA4 MTCE was effective against the TT cell line, it was totally ineffective in the MZ cell line, as cytotoxicity was driven entirely by the addition of TKIs. This mirrors a situation where a single MTCE is used and tumors show resistance via antigen loss, highlighting the importance of targeting multiple antigens simultaneously.

**Figure 10:**
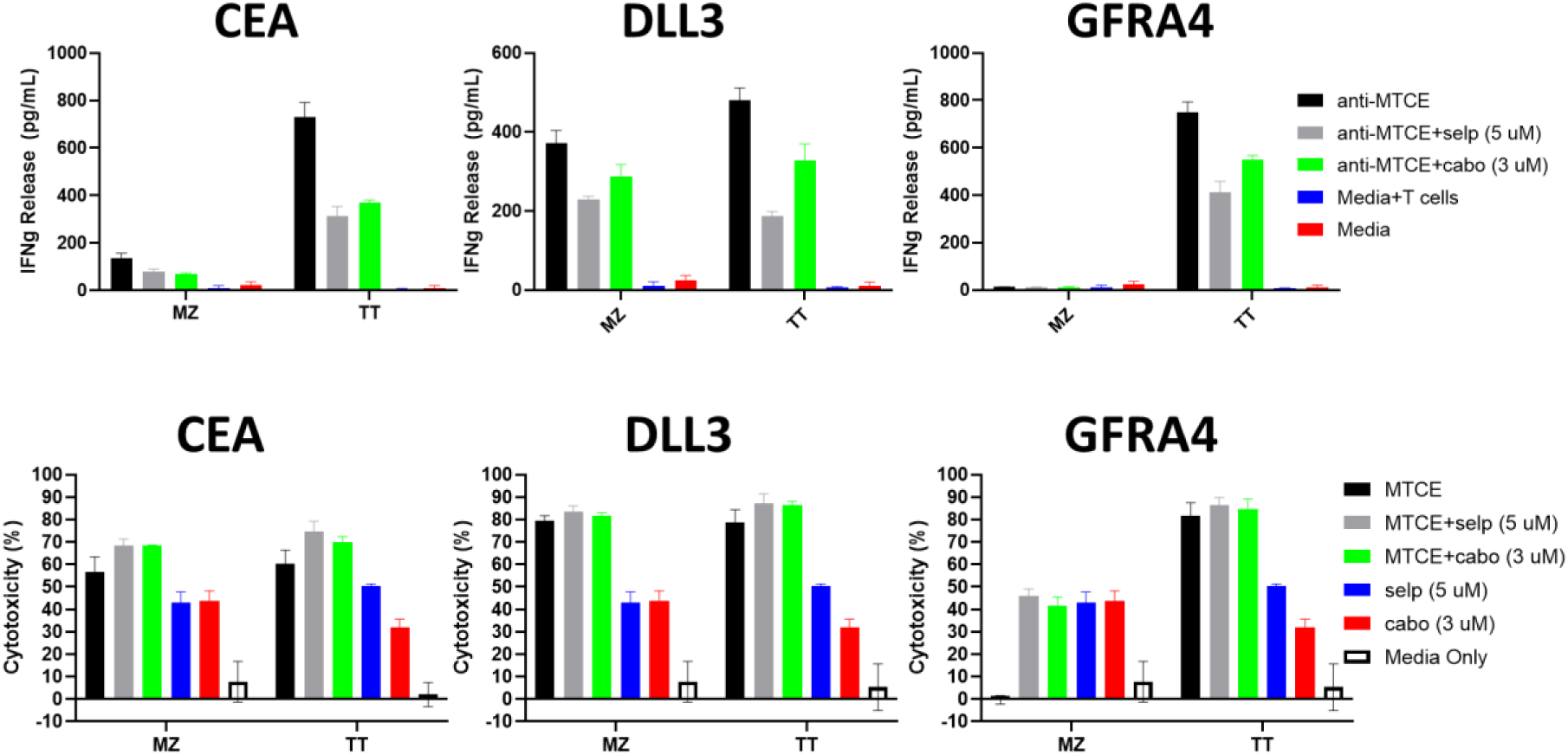
Effect of Combining TKIs with MTCEs. TKIs moderate IFNγ secretion, while numerically increasing cytotoxicity.

### Combining Multiple MTCEs

The anti-GFRA4 resistant MZ cells were used as an MTC cell line model system of tumor resistance mediated by target-antigen loss. As a countermeasure against antigen escape, all three MTCEs were combined at a concentration of 333 ng/mL each vs. individual MTCEs at a concentration of 1 µg/mL. The concentration was adjusted to keep the total amount of MTCEs in the cell culture constant. For the experiment, 15,000 MZ cells were plated overnight and then co-cultured for 24 hours with 7,500 T cells in the presence of MTCEs as indicated. Cytotoxicity was measured via XTT assay. As shown in Figure 11A, the combination of all three MTCEs at reduced doses mediated potent cytotoxicity and could potentially overcome tumor escape mediated by loss of a single target antigen (GFRA4). Interestingly, the triple combo yielded increased cytotoxicity over anti-CEA alone, which was likely driven by anti-DLL3, consistent with prior experiments showing that anti-DLL3 is the most potent MTCE against MZ cells. Triple MTCE treatment at reduced doses yielded comparable cytotoxicity to anti-DLL3 monotherapy at a higher dose, highlighting the feasibility of simultaneously targeting all three antigens to prevent antigen-loss escape.

**Figure 11:**
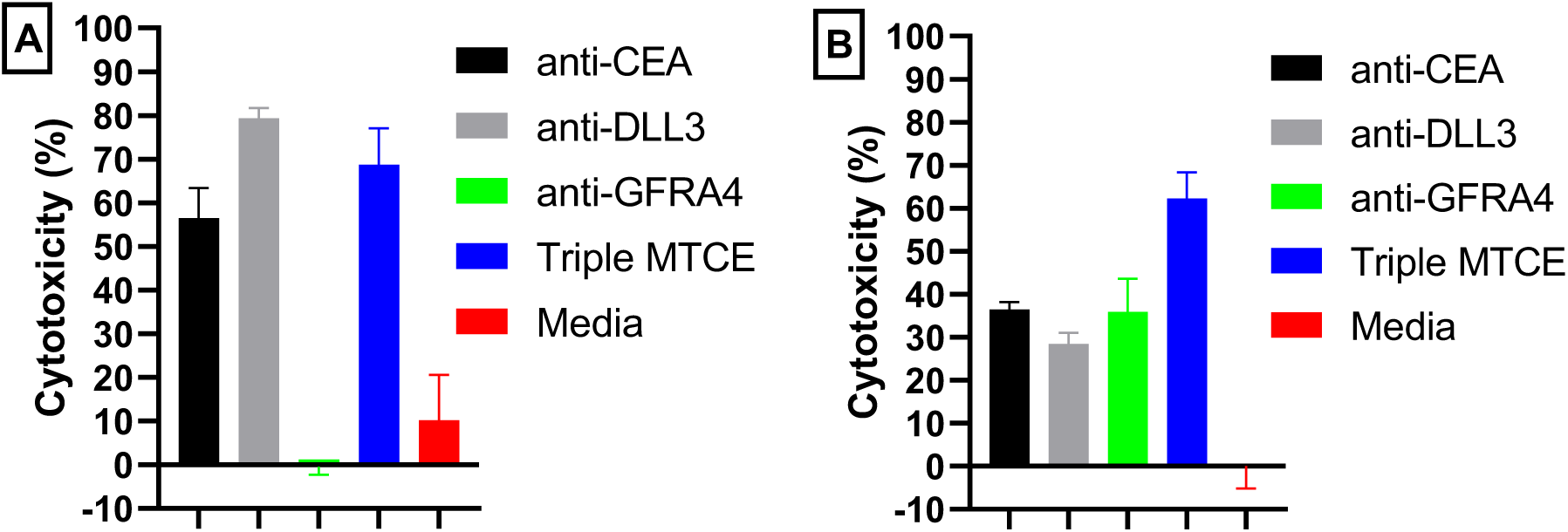
Combining MTCEs to Prevent Single Antigen Loss Escape. MTCEs can be combined to prevent antigen-loss escape in the anti-GFRA4 resistant MZ cell line (A). In a highly heterogeneous tumor comprised of CHO-CEA, CHO-DLL3, and CHO-GFRA4 cells, combining all three MTCEs significantly increased cytotoxicity over monotherapy (B).

To further assess the ability of reduced dose combination MTCEs to prevent or mitigate single antigen-loss escape, a highly heterogenous tumor was modeled by plating 5,000 CHO-CEA, 5,000 CHO-DLL3 and 5,000 CHO-GFRA4 cells (15,000 total) into each well. 7,500 T cells were then added to each well with 1 µg/mL of each MTCE individually or a combination off all three MTCEs at a concentration of 333 ng/mL. Cells were then cultured for 18 hours, and then cytotoxicity was measured by XTT assay. Despite reducing the concentration of each MTCE by 1/3, the Triple MTCE group mediated a significant increase (p < 0.001) in cytotoxicity over all monotherapy groups (Figure 11B), validating the strategy of combining multiple MTCEs to mitigate antigen loss escape in heterogenous tumors.

### Assessing MTCE Mechanism of Action in Multiple Biological Donors

Notably, all experiments up to this point in the manuscript were conducted using T cells from a single donor. In order to show that the mechanism of action of MTCEs is generalizable, and not unique to one individual and to characterize biological variability, a 48-hour XTT cytotoxicity assay was conducted using CD8 T cells from five healthy donors. CD8 T cells were isolated and cultured as previously described. For the experiment, 15,000 T cells were co-cultured with 15,000 tumor cells (TT, MZ, LNCaP (negative control #1), CHO-wt (negative control #2) CHO-CEA, CHO-DLL3 or CHO-GFRA4 cells) in the presence of 10 µg/mL of anti-CEA MTCE, anti-DLL3 MTCE (tarlatamab), anti-GFRA4 MTCE, gresonitamab (anti-CLDN18.2) or media only. The data is shown in Figure 12, and each data point represents the triplicate average from each unique donor. While there is clear variability in cytotoxicity, potentially related to intrinsic T cell fitness, all MTCEs mediate cytotoxicity in a target-dependent manner consistent with prior experiments. Notably, no cytotoxicity is observed in the target-antigen-negative CHO-wt or LNCaP cell lines, and the anti-CLDN18.2 TCE does not mediate cytotoxicity against either MTC cell line. The anti-CEA MTCE mediates strong cytotoxicity against TT cells and CHO-CEA cells and mediated moderate cytotoxicity against MZ cells, whereas tarlatamab is effective in both MTC cells lines and CHO-DLL3 only. Further, anti-GFRA4 only mediates strong activity against TT cells and CHO-GFRA4 cells. In summary, these data strongly indicate that the mechanism of action of MTCEs is consistent across human donors, and not unique to a single donor.

**Figure 12:**
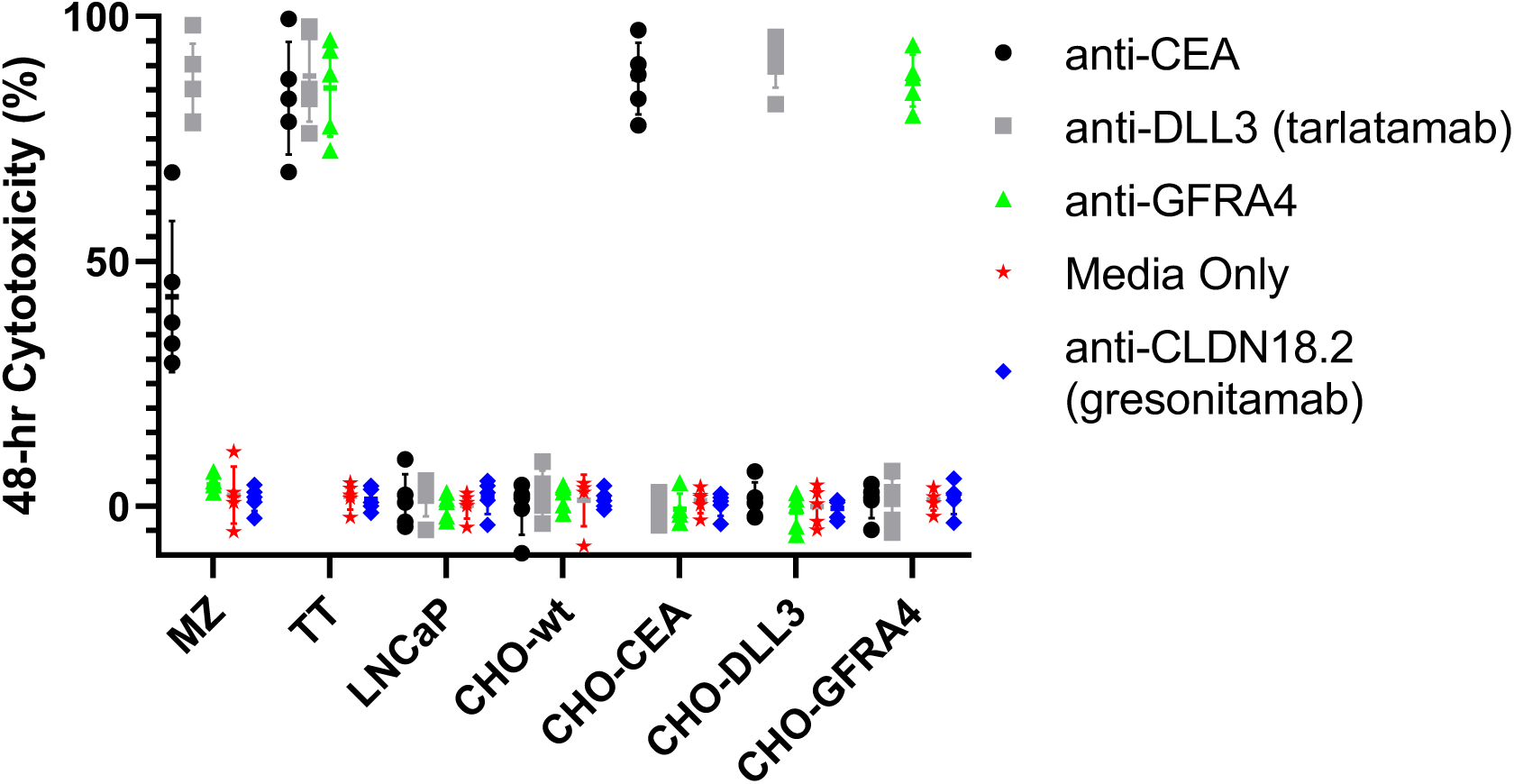
MTCEs mediate toxicity against tumor cell lines in a target-dependent manner in five independent healthy donors, with no observed cytotoxicity above baseline in the presence of gresonitamab or media only.

## Methods

### RNASeq and Gene Expression Data

MTC whole exome RNASeq data was acquired as previously described [19], and the entire dataset is fully disclosed in this manuscript for the first time. MTC patient paraffinized tumor specimens were collected and used to generate RNASeq data after Institutional Review Board (IRB) approval (IRB #16-1461), and all experiments were conducted in accordance with applicable regulations and guidelines under IRB approval. Further, written informed consent was obtained from all patients, with documentation on file at CU-Anschutz Medical Center (Aurora, CO, USA) [19]. Tumor cell line RNASeq data was downloaded from the TRON Cell Line Database (https://github.com/TRON-Bioinformatics/TCLP). Surfaceome genes were acquired from Table S2 in [20]. The Human Protein Atlas online portal (https://www.proteinatlas.org/) was used to evaluate protein expression and distribution in normal tissues. Analysis was performed using Excel software and all graphs were generated using GraphPad Prism software.

### Synthesis of anti-CEA and anti-GFRA4 MTCEs

MTCEs were synthesized at GenScript using their proprietary TurboCHO^TM^ transient transfection and purification process. Protein sequences including a designated signal peptide were disclosed to GenScript. The protein sequence was then codon optimized, and the corresponding gene was then cloned into pcDNA3.4 plasmids. After plasmid amplification and purification, CHO-S cells were transiently transfected with plasmids encoding each MTCE. After 5 days, the supernatant containing each MTCE was harvested and purified using Protein A chromatography followed by size exclusion chromatography followed by buffer exchange. Each MTCE was then molecularly characterized using SDS-PAGE and SEC-HPLC with both methods indicating purity exceeding 90% for anti-CEA MTCE and anti-GFRA4 MTCE. MTCE concentration was measured using A280 absorbance spectroscopy.

### Tumor Cell Lines, T cells and Cell Culture

With the exception of the MZ cell line, all cell lines were purchased from ATCC. The MZ cell was kindly received as a gift from Dr. Jena French (UC-Denver). Tumor cell lines were generally cultured in RPMI (CHO, Raji, LNCaP), DMEM (MZ, SKOV3, SHP-77) or Ham’s F12K (TT) media supplemented with 10% FCS and antibiotics. Cells were maintained with Normocin (InvivoGen) to prevent fungal and mycoplasma contamination. CHO-CEA, CHO-DLL3 and CHO-GFRA4 were generated by transducing CHO cells with lentiviral vectors encoding each target gene under control of an EF1-a promoter (Vectorbuilder). Cells were transduced with polybrene (8 µg/mL, MOI=10) before single cell cloning through limiting dilution, in order to establish pure monoclonal population of 100% transduced cells.

T cells used in single-donor experiments were generated as follows. First, the author’s peripheral blood PBMCs were isolated from whole blood using Ficoll-Paque centrifugation. Then cells were cultured in T cell media (AIM-V supplemented with 5% FCS, antibiotics and 100 IU/mL of IL-2) for 2 days in GREX 24-well plates at a concentration of 2e6 cells in 4 mL of media. After 2 days, CD8 T cells or bulk T cells were isolated using “untouched” commercial negative selection kits (#130-096-495, #130-096-535) from Miltenyi Biotech, prior to being used in assays and T cell purity exceeding 97% was confirmed by flow cytometry. For the experiment used to assess biological variability, healthy donor PBCMs were purchased from StemCell and processed in a manner identical to single donor experiments.

### Flow Cytometry

Flow staining antibodies were acquired from commercial vendors, as indicated in Table 2. For surface antigen staining of tumor cell lines, cells were gently detached using Accutase for 5-10 minutes, in order to avoid Trypsin-mediated protein cleavage [40]. Then cells were neutralized using ice-cold cell culture media. Cells were then spun down and resuspended in ice-cold media prior to staining 200,000 cells in 200 uL of media for 30 minutes at 4C. Staining antibodies were generally used at 1:100 dilution. After staining, cells were washed twice in cold FACS buffer, and then immediately analyzed on a BD Accuri flow cytometer.

**Table 2:** Antibodies used in experiments.

| Antibody | Clone | Supplier | Catalog Number |
| --- | --- | --- | --- |
| CD8-FITC | HIT8a | Biolegend | 500906 |
| CD4-FITC | OKT4 | Sony | 2187040 |
| Human IgG-PE | N/A | Jackson ImmunoResearch | 109-116-170 |
| CD3-PerCp | UCHT1 | Biolegend | 300428 |
| CEA-APC | CI-P83-1 | Santa Cruz Bio | sc-23928 |
| DLL3-PE | tarlatamab | MedChemExpress | HY-P99575 |
| GFRA4-PE | polyclonal | Saphire Bioscience | BS-0168R-PE |
| CD20-FITC | 2H7 | Biolegend | 302203 |
| CD16-FITC | 3G8 | Biolegend | 980112 |
| CD56-APC | QA17A16 | Biolegend | 985906 |
| CD14-FITC | M5E2 | Biolegend | 982502 |

### IFNγ Measurement

IFNγ was measured using an ELISA kit from Biolegend (#430101) in accordance with the manufacturer’s instructions. Prior to measurement, samples were diluted 1:5 in assay buffer, and this dilution factor was then corrected in the final results. ELISA plates were read on a Biotek EL808 plate reader at 450 and 650 nm.

### Cell Co-culture and XTT Cytotoxicity Assays

For each assay, tumor cells and T cells were cultured in a 50/50 blend of T cell media and tumor cell media in the presence of MTCEs as indicated. Wells were run in triplicate. To mitigate cytotoxicity from alloreactivity, each well was supplemented with the anti-MHC Class I blocking antibody W6/32 (Biolegend) at a concentration of 40 µg/mL. After 24 or 48 hours, the plate was flicked to remove non-adherent cells (dead cells or T cells). The plate was then washed once with warm media, flicked again and then 100 uL of media was added to each well, after which 50 uL of XTT reagent was added to each well, in accordance with instructions provided in the assay kit (Biotium, #30007). After XTT reagent addition, cells were then cultured for an additional 8 hours. Then plates were read on a Biotek EL808 ELISA plate reader at 490 nm and 650 nm. For analysis, the background was subtracted from each well (A_490_ – A_650_) and then denoted as A_sample_. Then MTCE-mediated cytotoxicity was computed as follows. The T cells + media only condition (A_baseline_) was used as a baseline. The MTCE-specific cytotoxicity was calculated as [(A_baseline_-A_sample_)/A_baseline_] x 100%. Under this calculation, when sample and baseline wells show equivalent absorbance (A_baseline_ = A_sample_) due to wells harboring the same number of metabolically active tumor cells, cytotoxicity is zero. Conversely, when A_sample_ is zero due to elimination of all metabolically active cells, then cytotoxicity is 100%.

## Discussion

The application of T cell engagers in MTC is a novel therapeutic strategy. MTCEs enable endogenous T cells to robustly lyse MTC cells expressing the targeted antigens CEA, DLL3 or GFRA4, *in vitro*. These antigens were identified by leveraging data from an MTC patient dataset in conjunction with differential gene expression analysis of large cell line database, and their expression was confirmed on MTC cell lines. While anti-CEA and anti-GFRA4 MTCEs are novel molecular entities, tarlatamab (anti-DLL3) is already FDA approved. Importantly, both anti-CEA and anti-GFRA exhibited robust *in vitro* activity at doses (∼10 ng/mL) well below the average and trough concentrations of tarlatamab measured in clinical trials (C_avg_ = 1040 ng/mL and C_trough_ = 495 ng/mL) [40]. In terms of monotherapy, tarlatamab consistently generated the most consistent cytotoxicity across both cell lines, even at low concentrations, and is thus immediately recommended for investigational off-label treatment in MTC patients whose tumors highly express DLL3.

While tarlatamab has demonstrated robust clinical activity, it is notable that the reported complete response rate in SCLC is only 2% [29]. As tumor heterogeneity and plasticity are hallmarks of solid tumors [42–44], it is likely that the rarity of complete responses could be due to loss or downregulation of DLL3 expression. Targeting more than a single antigen may be necessary for tumor eradication in the majority of MTC patients, and bulk tumor transcriptomic data highlights variable expression of all three antigens (Figure S3). Thus, the utilization of MTCEs targeting three independent antigens (CEA, DLL3, GFRA4) represents a compelling strategy to overcome antigen-loss escape [45]. Recent scRNA analysis has demonstrated that multiple MTC subtypes with distinct transcriptional programs can exist within the same tumor, indicating that MTC cells can exist in multiple states during the time-course of evolution from C cell, to differentiated tumor cells, to dedifferentiated and aggressive malignant cells [44]. The transcriptional changes can influence antigen expression, as observed in the MZ cell line, which shows minimal GFRA4 expression, moderate CEA expression and high DLL3 expression. However, it is notable that both MTC cell lines express a triple-negative population by flow, which could lead to tumor recurrence even under optimal MTCE therapy. It is unclear if antigen-expression is cell-cycle dependent, or if cells may temporally recover expression at later timepoints. Further analysis of this population by flow sorting, followed by re-culturing and epigenetic methylation analysis would be particularly illuminating.

A prerequisite for MTCEs to function is the presence of T cells in close proximity to tumor cells. While MTC is generally thought to be a cold tumor lacking significant T cell infiltration, T cells have been found in the majority of MTC primary tumors although data on distant metastases is lacking. However, T cell engagers are active in other immunologically cold tumor types, such as prostate cancer, indicating that T cells may exist in sufficient quantities to generate an antitumor response. Notably, the lymph nodes are often the first observable site of metastases in MTC [46], and therefore lymph node tumors may be particularly amenable to MTCE therapy, as the tumor would inherently be in close proximity to T cells. Tertiary lymphoid structures could be another source of T cells, and it is possible that initial killing by sparse T cells could lead to inflammation and trafficking of additional T cells to the tumor site. Notably, MTCEs were observed to be functionally combinable with TKIs, which can relieve immunosuppression in the TME through a variety of mechanisms. These T cells could further be supported by costimulatory agonist antibodies and/or novel IL-2 analogs, in order to overcome the immunosuppressive TME. As MTC is a rare tumor type, there is still an incomplete picture on the immune composition at common distant metastatic sites, including the lungs, liver and bones. Data on both the immune infiltrate and target antigen expression in distant metastases would be particularly helpful in identifying patients who could respond to MTCE therapy. However, the observation that higher tumor burden correlates with higher serum CEA indicates that many distant metastases retain CEA expression.

Our data indicates a more favorable ability to eradicate tumor cells at higher effector to target ratios. In the clinical setting, it is therefore likely that patients with minimal residual disease would have higher response and greater chances of a cure than patients with bulky metastatic disease. However, clinical data would ultimately be needed to fully test this hypothesis. As there is no effective adjuvant therapy for MTC, MTCEs might represent a compelling treatment to eradicate residual metastatic disease and ultimately cure patients who would otherwise recur. This would be analogous to radioactive iodine therapy in differentiated thyroid cancer patients. For patients with bulky metastatic disease, our data indicates that MTCEs are functionally compatible with standard-of-care TKIs, and that the two treatment modalities may be synergistic in the clinic. Reduced tumor burden would correspond to fewer activated T cells, and therefore less cytokine release, which is a dose-limiting toxicity of T cell engagers. Additionally, combination therapy may further shift the equilibrium toward tumor eradication over tumor growth.

The proof-of-concept data presented herein demonstrate MTCEs as a novel off-the-shelf strategy for redirecting the immune system to eradicate MTC. While *in silico* immunogenicity prediction results generated using the AbImmPred and CD4 Episcore algorithms (Tables S1 and S2) predicted immunogenicity levels in-line with FDA-approved antibodies [47,48], additional preclinical *in vitro* experiments to assess potential immunogenicity are warranted [49–50]. Additional assays to more fully assess any potential off-target binding of MTCEs will be conducted. Characterization of target expression levels and heterogeneity in distant metastases along with a comprehensive analysis of the tumor-immune microenvironment would be helpful in identifying patients who are likely to benefit from MTCE therapy. These studies will be the subject of future efforts, with the ultimate aim of bringing multi-antigen-targeting MTCEs into the clinic and radically improving the treatment of metastatic MTC.

## Acknowledgements

The author would like to personally thank Dr. Jena French (CU-Anschutz Medical Center) for many helpful discussions and note that the MTC patient RNASeq data extensively analyzed for MTCE development *de novo* in this manuscript was generated during a prior study and was the subject of the publication titled, “Comprehensive immune profiling of medullary thyroid cancer” which is reference [19]. The author would also like to thank Dr. Eric Tran (Providence) for countless helpful discussions, being a wonderful mentor and allowing use of his lab’s fluorescence microscope for imaging.

## Funding

This research was solely funded by ThyThera, LLC.

## Author Contributions

This manuscript was written solely by Tim Erickson, and all experiments were designed and conducted by Tim Erickson.

---

**Figure S1.**
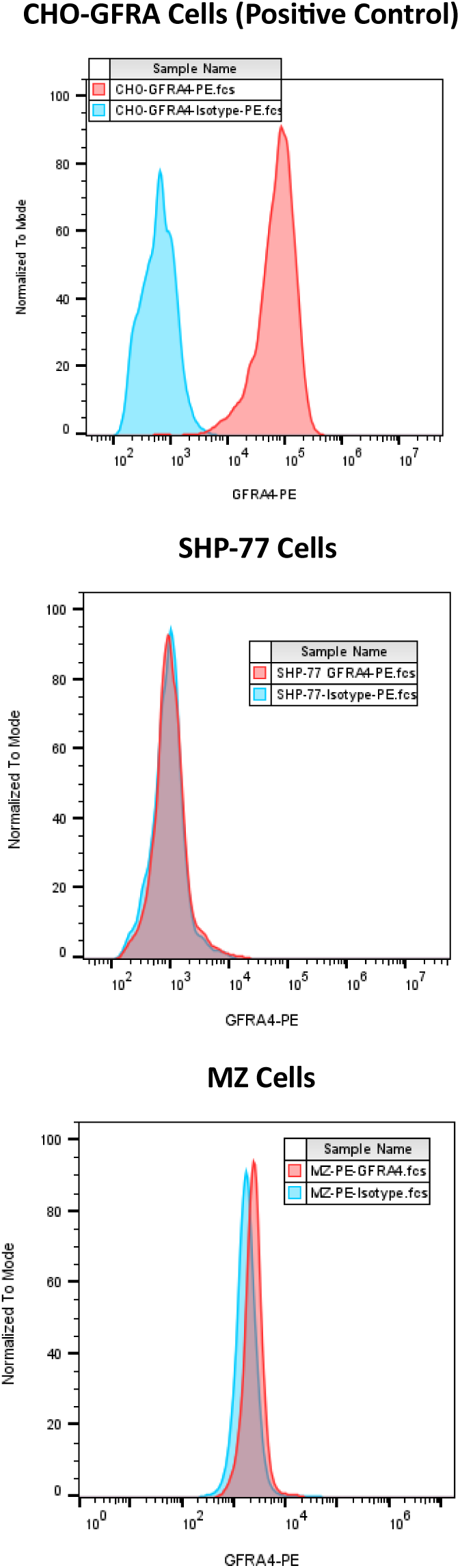
Single-stain (PE only) detection of GFRA4 on CHO-GFRA4, SHP-77 and MZ cells. Relative to isotype control, there is no detectable GFRA4 expression on SHP-77 cells, whereas there is very dim, but detectable expression of GFRA4 on MZ cells.

**Figure S2:**
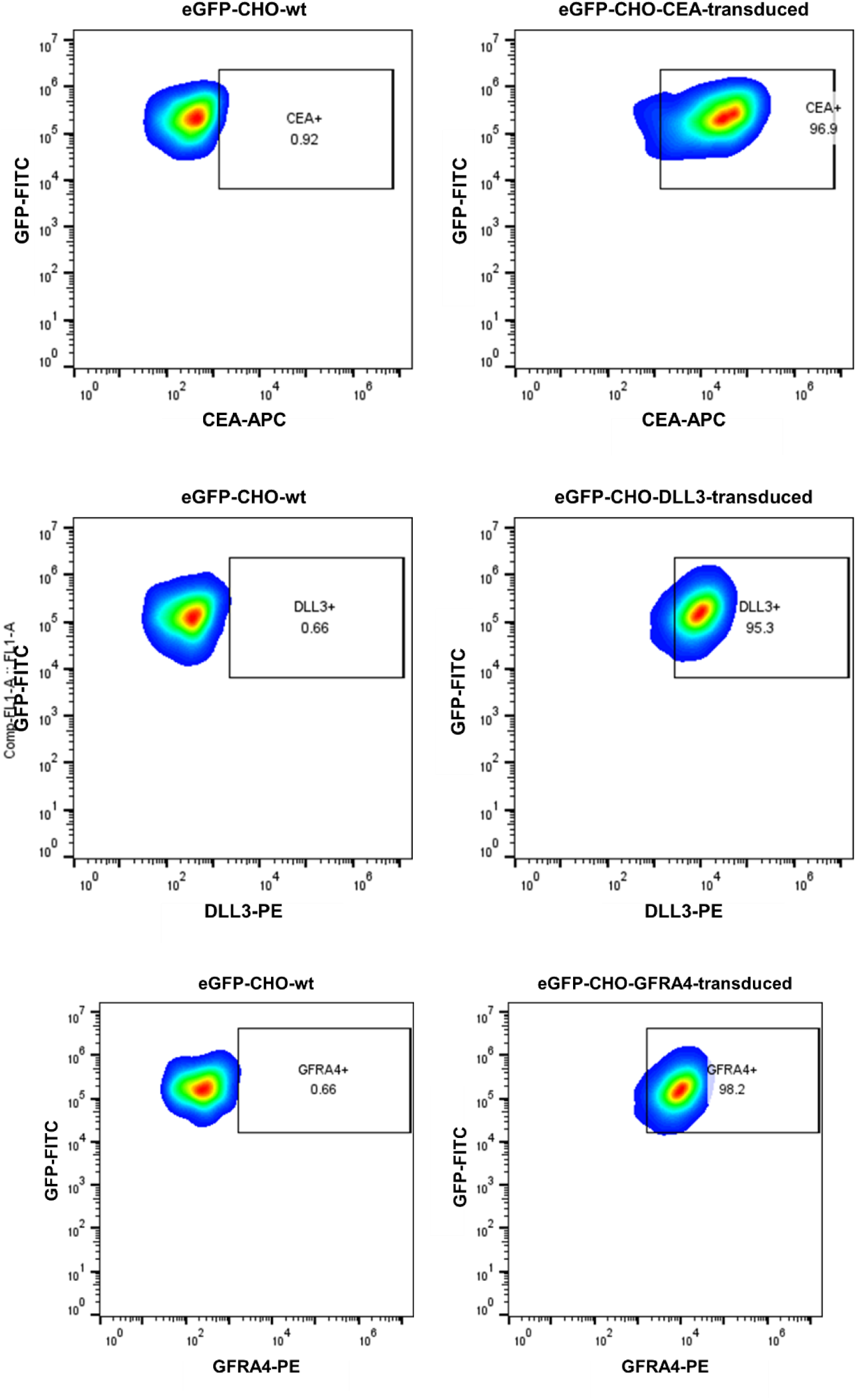
Flow confirmation of surface target antigen expression on target-transduced CHO cells. Cells were generated from a clonal population of GFP-transduced CHO cells, in order to facilitate target cell identification in live-cell imaging studies.

**Figure S3:**
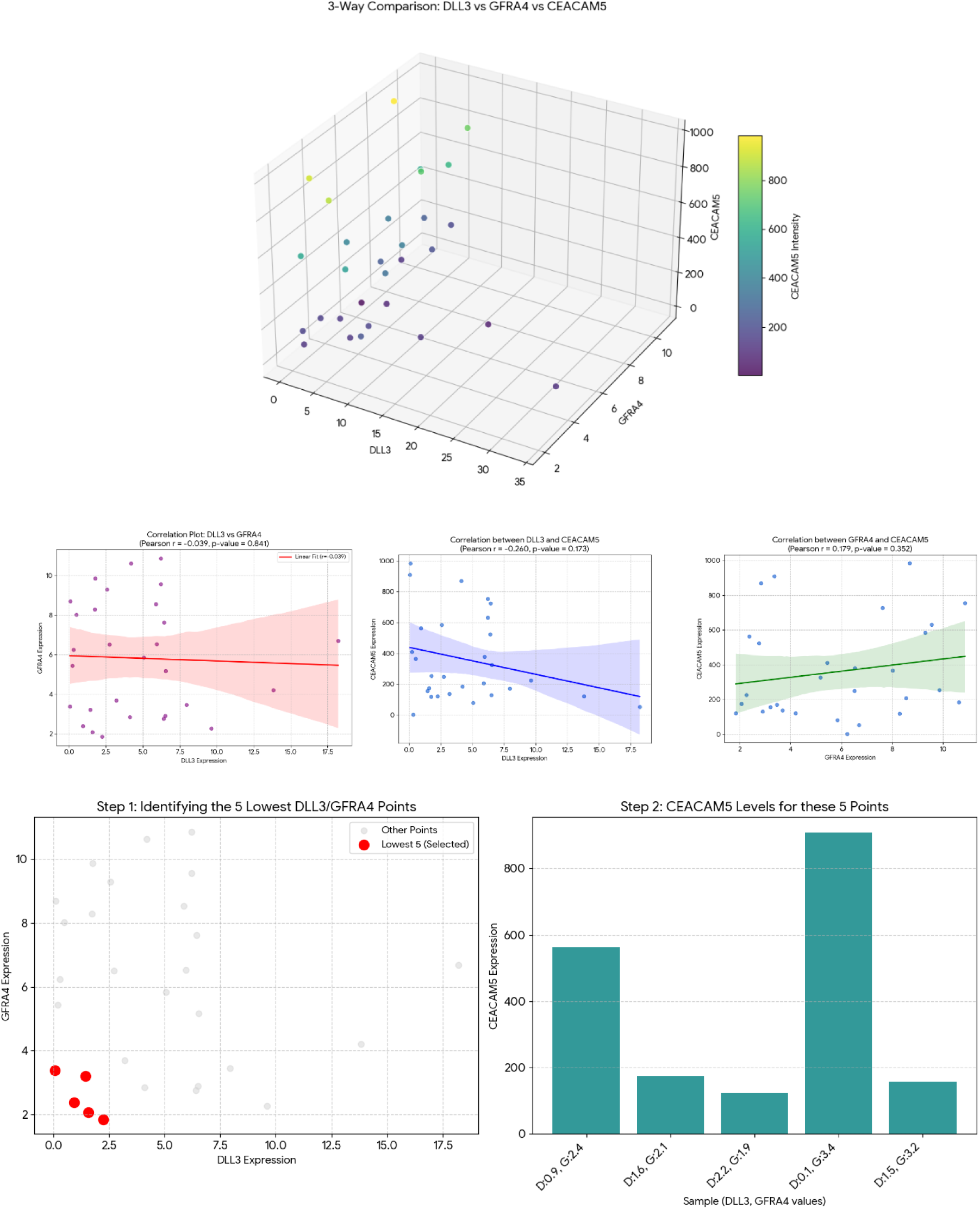
Transcriptomic data (units=FPKM) from MTC tumors (n=30) shows heterogeneity in CEA, DLL3 and GFRA4 expression (top) with no definitive statistical correlation observed among these antigens (middle). Notably all patients with low DLL3 and low GFRA4 expression (red dots) had high levels of tumoral CEA expression. A single patient with undetectable DLL3 expression had observable GFRA4 expression.

**Table S1:**
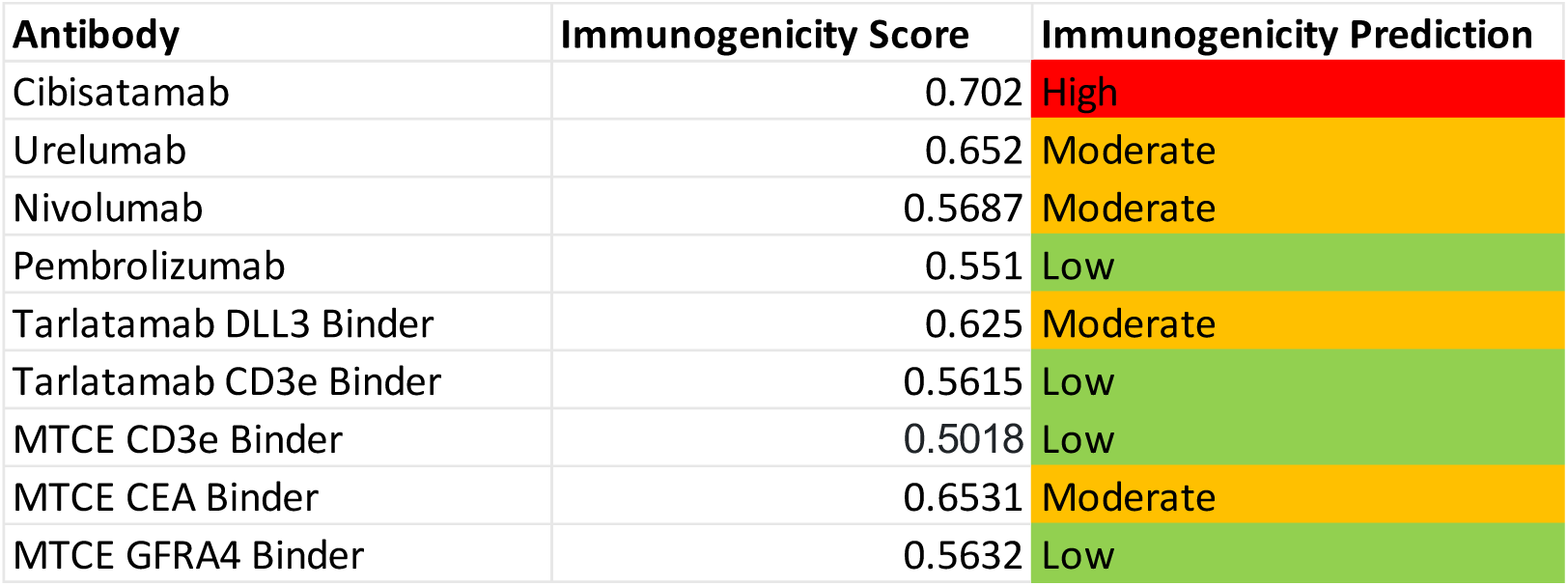
AbImmPred Immunogenicity Prediction Results [49]. Both the MTCE CD3e binder and GFRA4 binder are predicted to have low immunogenicity. The CEA binder is predicted to have moderate immunogenicity, but is only slightly higher than FDA-approved tarlatamab and comparable to urelumab.

| Antibody | Immunogenicity Score | Immunogenicity Prediction |
| --- | --- | --- |
| Cibisatamab | 0.702 | High |
| Urelumab | 0.652 | Moderate |
| Nivolumab | 0.5687 | Moderate |
| Pembrolizumab | 0.551 | Low |
| Tarlatamab DLL3 Binder | 0.625 | Moderate |
| Tarlatamab CD3e Binder | 0.5615 | Low |
| MTCE CD3e Binder | 0.5018 | Low |
| MTCE CEA Binder | 0.6531 | Moderate |
| MTCE GFRA4 Binder | 0.5632 | Low |

**Table S2:**
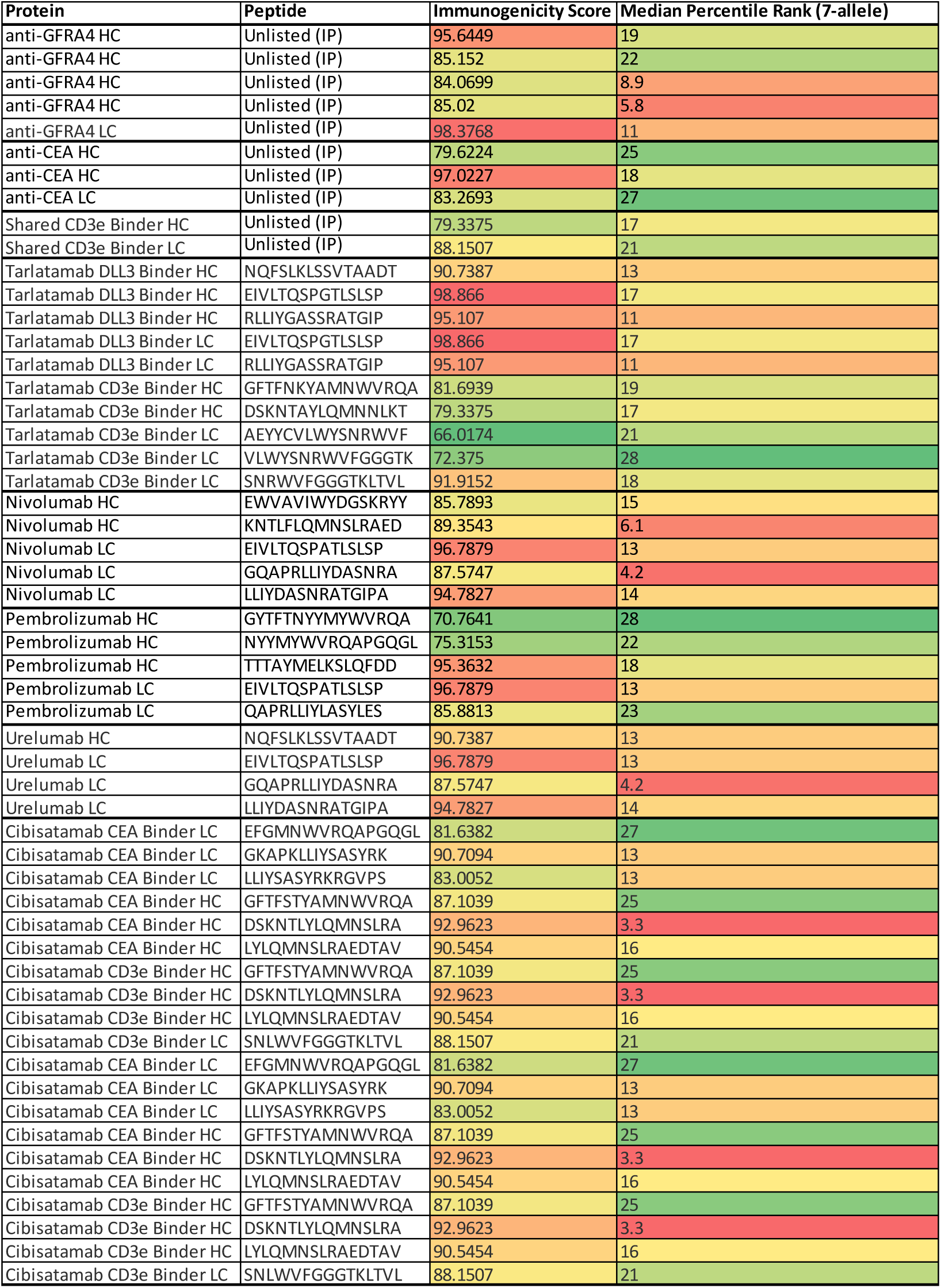
Prediction of CD4 epitopes using the CD4 Episcore algorithm [50]. Both the number of predicted epitopes for each construct and immunogenicity score are shown. Note the high number of predicted CD4 epitopes for cibisatamab, which was withdrawn from clinical development due to immunogenicity concerns. Including the CD3e binder, anti-GFRA4 harbors seven predicted epitopes, whereas anti-CEA harbors five. This is similar to nivolumab (five), pembrolizumab (five) and urelumab (four), and less than tarlatamab (ten) and much less than cibisatamab (twenty). Cibisatamab elicited broad neutralizing antibody responses in the majority of patients, and development was abandoned by Roche.

## Notes

### Competing Interest Statement

The authors have declared no competing interest.

### Summary of Updates

Additional figures and text have been added in response to reviewer comments.

